# AAV and lentiviral transduction in Duchenne muscular dystrophy cardiomyocytes activate cell stress responses

**DOI:** 10.64898/2026.07.31.742163

**Authors:** Elaine C. Lai, Abiageal R. Keegan, Asuka Eguchi

**Author notes:** **Correspondence should be addressed to:** Asuka Eguchi, 845 Health Sciences Road, Gross Hall 4052, Irvine, CA 92697.

## Abstract

Duchenne muscular dystrophy (DMD) is an X-linked muscle wasting disorder marked by lack of dystrophin expression. Symptoms include loss of ambulation, respiratory problems, and cardiac complications with heart failure being the leading cause of death. Dystrophin transduces force from the actin cytoskeleton to the extracellular matrix to protect cells during muscle contraction. Restoration of dystrophin expression by gene transfer holds promise in addressing the root cause of disease. We compared the changes to transcriptional profiles after gene transfer by adeno-associated virus or lentivirus to examine whether viral treatment alone impacts cell homeostasis. We delivered GFP to cardiomyocytes differentiated from induced pluripotent stem cells (iPSCs) with DMD mutations. Global transcriptional profiling revealed a downregulation of metabolic genes after lentiviral transduction compared to untreated controls. In both AAV and lentivirus-treated DMD iPSC-cardiomyocytes, we observed an activation of the p53 DNA damage response in addition to a downregulation of cell cycle genes, suggesting stress-induced G2/M checkpoint arrest following viral delivery. These findings demonstrate that gene therapy mediated by viral vectors activates cell stress pathways. Interventions to mitigate these stress responses may be necessary for safe and effective gene transfer in diseased cells.

## INTRODUCTION

Duchenne muscular dystrophy (DMD) is an X-linked progressive muscle wasting disorder affecting approximately 1 in 5,000 male births^1^. DMD is caused by mutations in dystrophin, resulting in an absence of dystrophin protein. Dystrophin is a critical structural component of the dystrophin-associated protein complex, spanning the actin cytoskeleton to sarcolemma^2^. In muscle fibers, dystrophin transduces the forces generated during muscle contractions and distributes the force to the extracellular matrix to protect the plasma membrane from mechanical stress. Without dystrophin, repeated cycles of contraction lead to progressive membrane damage, chronic inflammation, and replacement of muscle tissue with fibrotic and adipose tissue^3^.

The clinical course of DMD is characterized by progressive skeletal muscle weakness, with patients losing ambulation, experiencing respiratory problems, and manifesting cardiac complications. Heart failure is now the leading cause of death for patients in their third decade of life^4^. The cardiac issues in DMD is particularly significant because the heart has little capacity to regenerate and thus cannot replace the cardiomyocytes lost due to contraction-mediated injury. Consequently, the heart is one of the most challenging, critical organ targets for DMD therapeutics.

Viral gene therapy has emerged as one of the most promising strategies to address the root cause of DMD, the lack of the dystrophin protein. Adeno-associated virus (AAV) is currently the most efficient way to systemically deliver genes to the skeletal muscle, diaphragm and heart. However, systemic administration of AAV vectors at the high doses required to achieve therapeutic levels of transduction throughout the skeletal muscles in the body has been associated with serious adverse events in both preclinical models and clinical trials^6–10^. These include fatal immune-mediated thrombotic microangiopathy, acute liver failure, and myocarditis. Investigations into these outcomes have highlighted the role of innate and adaptive immune responses to the AAV capsid and transgene product^10^. The contribution of cell-intrinsic stress responses to AAV-mediated toxicity has not been well characterized.

In neurons differentiated from human induced pluripotent stem cells (iPSCs), transduction with AAV delivering GFP induced an inflammatory gene signature and activation of p53-dependent DNA damage response pathways^11^. These observations suggest that the stress imposed by viral delivery may represent another contributor to the severe adverse events observed in patients that have received AAV gene therapy. Understanding these responses is particularly important in the context of diseased cells, which may be more vulnerable to additional cellular stressors.

Human iPSC-derived cardiomyocytes (iPSC-CMs) offer a platform to study the safety of gene therapy approaches. Previously, we have shown that DMD iPSC-CMs recapitulate key features of disease, including aberrant calcium handling, contractile dysfunction, and poor cell viability^12–14^. To investigate how viral gene delivery affects cellular homeostasis in the context of DMD, we delivered GFP with either AAV serotype 6 (AAV6) or a lentiviral vector to iPSC-CMs bearing a nonsense mutation in the dystrophin gene. We performed bulk RNA sequencing at two timepoints following transduction to capture both early and late transcriptional responses. Our results reveal that both AAV and lentivirus activate the p53 DNA damage response and suppress cell cycle progression genes, consistent with G2/M checkpoint arrest. Lentiviral transduction additionally suppresses a broad program of metabolic gene expression. Together, these findings identify cell stress activation as a common consequence of viral gene delivery in DMD cardiomyocytes and suggest that strategies to mitigate these responses may improve the safety and tolerability of viral gene therapy.

## RESULTS

### AAV and lentivirus treatment alter the transcriptional profile of DMD iPSC-cardiomyocytes

To determine whether viral gene delivery induces transcriptional changes in DMD cardiomyocytes, we transduced DMD iPSC-CMs with either AAV6 or lentiviral vectors encoding GFP under the control of the muscle-specific CK8 promoter. Transductions were performed at day 16 of differentiation, and cells were collected for bulk RNA sequencing at day 20 (4 days post-transduction) and day 30 (14 days post-transduction) to capture both acute and more sustained transcriptional responses.

Principal component analysis (PCA) of RNA sequencing data showed clear separation between the timepoints of untreated and virus-treated samples along the first principal component (PC1, 30% variance), while the second principal component (PC2, 10% variance) captured variability associated with treatment (**Figure 1**). Notably, AAV- and lentivirus-treated samples clustered together and apart from untreated controls, indicating that viral transduction drives a distinct and common transcriptional signatures regardless of vector type or time post-transduction.

**Figure 1.**
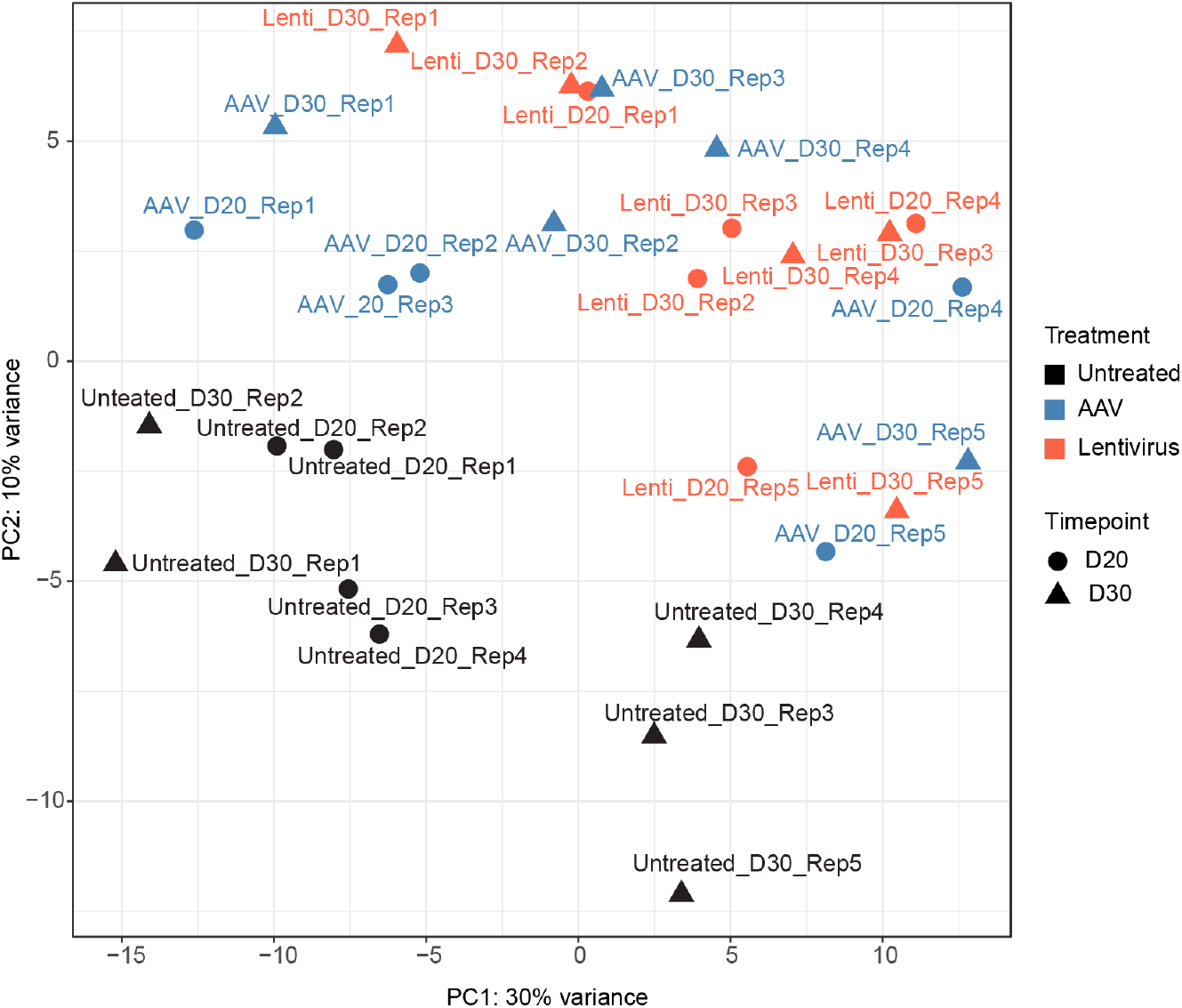
Principal component analysis of untreated, and AAV and lentivirus DMD iPSC-cardiomyocytes. D20=day 20, D30=day 30 of differentiation. Rep=replicate.

To further characterize the transcriptional changes induced by viral treatment, we performed differential gene expression analysis on all pairwise comparisons among treatment groups and timepoints. Differentially expressed genes (DEGs) were defined using a cutoff of absolute log_2_ fold change > 1 and a false discovery rate (FDR) < 0.05. Unsupervised hierarchical clustering of DEGs revealed five distinct gene expression clusters (**Figure 2A**). Cluster 1 was enriched for metabolic genes and exhibited higher expression in untreated cells that diminished in lentivirus-treated samples. Clusters 3 and 4 contained genes associated with cardiomyocyte differentiation. Cluster 2 contained genes related to differentiation and DNA damage signaling, while Cluster 5 was enriched for cell cycle progression genes and showed markedly lower expression in virus-treated cells at both timepoints. Taken together, the clustering patterns indicate that viral transduction suppresses programs of metabolic activity and cell cycle progression while activating stress response pathways.

**Figure 2.**
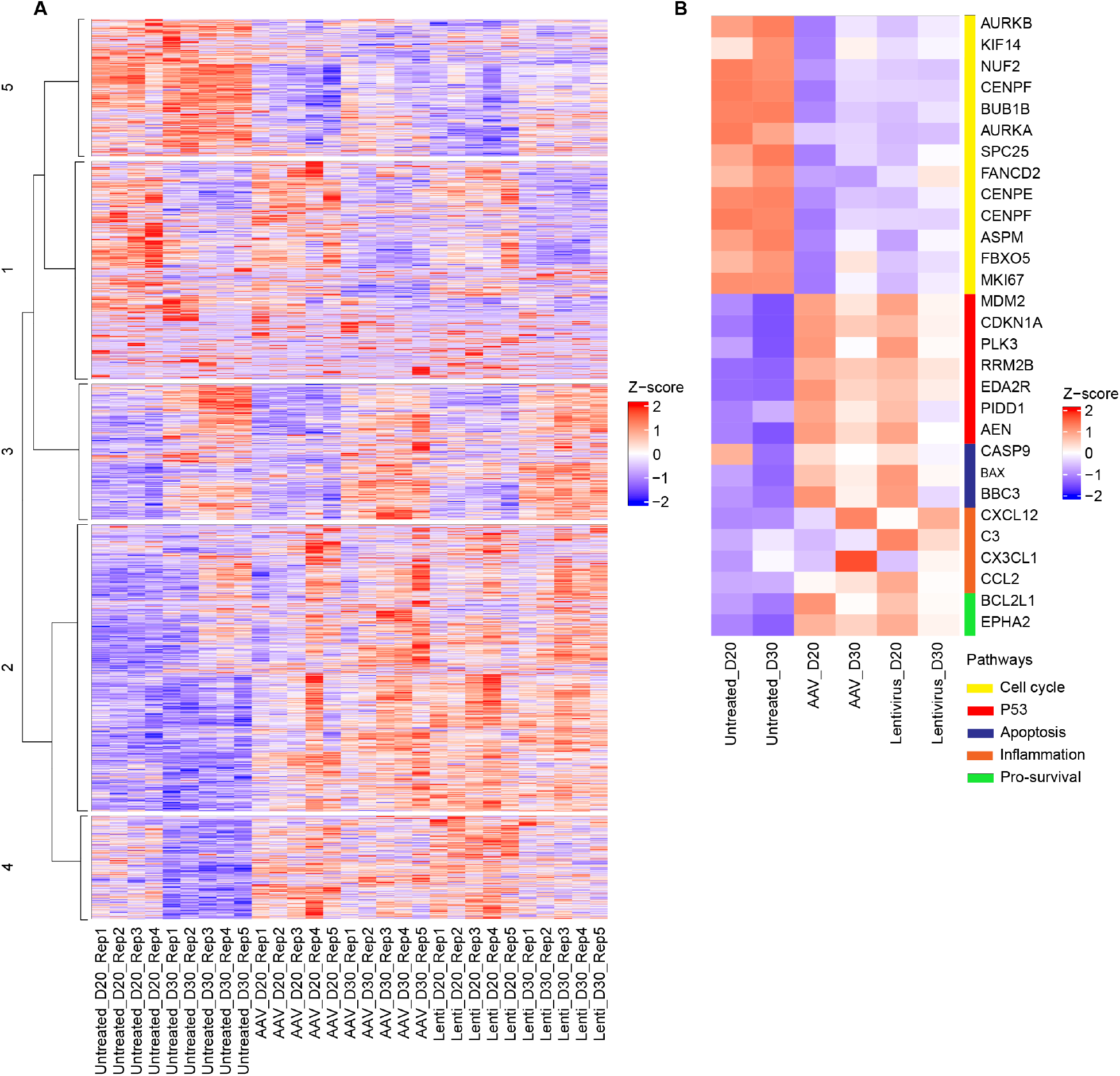
Transcriptional profile of AAV and lentivirus treated cells from each replicate. **A)** Heatmap of differentially expressed genes (DEGs) from each pairwise comparison. Unbiased heirchcial clustering revealed five clusters. DEGs were defined with a cut off value of absolute log_2_ fold change > 1 and FDR < 0.05. **B)** Heatmap of DEGs related to different pathways.

To more precisely define the pathways affected by viral treatment, we examined the expression of a subset of genes associated with five key biological processes: cell cycle regulation, p53 signaling, apoptosis, inflammation, and pro-survival responses (**Figure 2B**). Cell cycle genes critical for mitotic entry and progression, including Aurora kinase B (AURKB), Aurora kinase A (AURKA), KIF14, NUF2, CENPF, CENPE, BUB1B, SPC25, ASPM, FBXO5, and MKI67, were consistently downregulated in both AAV- and lentivirus-treated cells compared to untreated controls at both timepoints. This pattern of gene repression is consistent with arrest at the G2/M cell cycle checkpoint, in which cells halt division in response to DNA damage and stress prior to mitotic entry.

Multiple canonical p53 target genes were upregulated in virus-treated cells, including MDM2, CDKN1A (p21), PLK3, RRM2B, EDA2R, PIDD1, AEN, and FANCD2. CDKN1A encodes the cyclin-dependent kinase inhibitor p21, a direct p53 transcriptional target that enforces cell cycle arrest, and its upregulation is consistent with p53-mediated G2/M checkpoint activation. MDM2, another p53 transcriptional target and negative regulator of p53, was also upregulated, indicating a feedback response to contain p53 activity. The DNA repair gene FANCD2 was upregulated in virus-treated cells at both timepoints, further supporting the presence of DNA damage signaling.

Apoptotic pathway genes including CASP9, BAX, and BBC3 (PUMA) were upregulated in virus-treated samples. BBC3 and BAX are pro-apoptotic BCL-2 family members whose expression is induced by p53 in response to DNA damage, and CASP9 is a critical initiator of the intrinsic apoptotic cascade. Inflammatory chemokine and complement genes including CXCL12, C3, CX3CL1, and CCL2 were also upregulated in virus-treated cells, suggesting activation of innate immune-associated signaling even in the absence of immune cells.

Despite evidence of apoptotic gene induction, virus-treated cells also upregulated pro-survival factors BCL2L1 and EPHA2. BCL2L1 encodes Bcl-xL, an anti-apoptotic protein that inhibits mitochondrial outer membrane permeabilization and counteracts the pro-apoptotic functions of BAX and PUMA, suggesting that the virus treated DMD iPSC-CM exhibit a compensatory survival response following viral-induced stress. EPHA2, an Eph receptor tyrosine kinase with known roles in promoting cell survival and stress adaptation, may similarly reflect an attempt to sustain cellular viability under conditions of viral stress (cite). Collectively, these results indicate that both AAV and lentiviral transduction in DMD iPSC-CMs activate a coordinated stress response involving p53 signaling, DNA damage, G2/M checkpoint engagement, apoptosis, inflammation, and compensatory pro-survival signaling.

Comparison of AAV and lentivirus responses revealed both shared and distinct transcriptional features. While p53 activation and cell cycle suppression were common to both conditions, lentivirus-treated cells showed a more pronounced downregulation of Cluster 1 metabolic genes at both timepoints. This likely reflects genomic integration within open regions of chromatin which does not happen with AAV transduction as the AAV genome remains episomal.

## DISCUSSION

In this study, we conducted bulk RNA sequencing to examine the transcriptional consequences of AAV and lentiviral gene delivery in DMD iPSC-CMs. Our results demonstrate that viral transduction, independent of the therapeutic payload, activates a robust p53 DNA damage response and suppresses genes required for G2/M cell cycle progression. Lentiviral transduction additionally impairs the expression of metabolic genes. These findings extend previous observations from iPSC-derived neurons and establish that viral vector-mediated stress responses occur in cardiomyocytes derived from a disease-relevant DMD genetic background.

Our findings have direct translational implications for the design and clinical application of AAV gene therapy The observation that both AAV and lentiviral vectors activate p53 and suppress cell cycle genes suggests that these effects are not specific to one viral vector platform but represent a general feature of viral gene delivery. Strategies to reduce viral vector-associated cell stress may include optimization of vector dose and serotype or co-administration of anti-apoptotic activators or DNA damage inhibitors or at the time of gene delivery. The identification of BCL2L1 and EPHA2 as endogenously upregulated pro-survival factors in virus-treated cells also suggests that these pathways may represent targets for enhancing cardiomyocyte survival following gene transfer.

In summary, these findings underscore the importance of evaluating vector-intrinsic cellular effects in disease-relevant cell models and support the development of co-therapeutic strategies to improve the safety of AAV gene therapy in DMD and potentially other viral vectors for viral gene transfer.

## MATERIALS AND METHODS

### iPSCs

All human induced pluripotent stem cell lines were approved by and conducted in accordance with the Human Stem Cell Research Oversight Committee at the University of California, Irvine.

### Cardiomyocyte differentiation

iPSCs were seeded onto Matrigel (Fisher Scientific)-coated plates and maintained in StemFlex medium (Gibco). Upon reaching 60–70% confluency, cardiomyocyte differentiation was initiated by culturing cells in RPMI 1640 supplemented with 1× B27 minus insulin (Invitrogen) and 4 μM CHIR-99021 (Sigma-Aldrich) for 2 days. On day 2, medium was replaced with RPMI 1640 supplemented with 1× B27 minus insulin and 5 μM IWR-1 (Sigma-Aldrich) for an additional 2 days. On day 4, cells were transitioned to RPMI 1640 supplemented with 1× B27 (Invitrogen) and maintained for 4 days with medium changes every 2 days. On day 12 of differentiation, iPSC-derived cardiomyocytes (iPSC-CMs) were dissociated using Acutase and TrypLE and replated onto Matrigel-coated plates in RPMI 1640 minus glucose supplemented with 1× B27, 4 mM sodium lactate (Sigma-Aldrich), and 5% KnockOut Serum Replacement (KSR; Invitrogen). Medium was refreshed the following day and every 2 days thereafter with RPMI 1640 minus glucose supplemented with 1× B27 and 4 mM lactate. On day 16, iPSC-CMs were transduced with AAV6 at a multiplicity of infection at 30,0000 or lentiviral vectors encoding GFP and cultured until Day 20 or Day 30 of differentiation for downstream analyses.

### Cloning of viral vectors

The AAV transfer plasmid backbone was derived from pAAV-CAG-GFP (Addgene plasmid #37825), a gift from Edward Boyden. The muscle-specific CK8 promoter was PCR-amplified from a CK8-luciferase-zeocin plasmid. The 3’ UTR and synthetic poly-A signal were amplified from the H3μDys construct provided by Dr. Jeff Chamberlain (University of Washington). The final AAV transfer plasmid, pAAV-CK8-GFP, was assembled using Gibson Assembly by replacing the CAG promoter in pAAV-CAG-GFP with the CK8 promoter and appending the 3’ UTR and poly-A signal downstream of the GFP coding sequence.

### AAV production and purification

AAV production and purification protocol was adapted from Addgene. Recombinant AAV6 vectors were produced using a standard triple-plasmid transient transfection protocol in AAV293 cells. Briefly, HEK293T cells were seeded in 15 cm dishes and transfected at approximately 70– 80% confluency using Lipofectamine 3000. Cells were co-transfected with three plasmids: (1) the AAV transfer plasmid encoding CK8-GFP flanked by AAV2 inverted terminal repeats (ITRs); (2) the AAV rep and cap plasmid encoding the AAV2 rep genes and AAV6 cap gene (pRC6 or equivalent); and (3) the mini pHelper plasmid providing the adenoviral genes E2A, E4, and VA RNA required for AAV replication. Plasmids were combined at a molar ratio of 1:1:1 transfer:rep/cap:helper. At 48-72 hours post-transfection, cells were harvested by scraping and pelleted by centrifugation. Cells were lysed by sonication in lysis buffer and treated with benzonase nuclease (50 U/mL; Sigma-Aldrich) at 37°C for 30 minutes to degrade unpackaged nucleic acids. Cell debris was removed by centrifugation at 3,000 × g for 10 minutes at 4C.

AAV vectors were purified by ultracentrifugation using an iodixanol (OptiPrep) density gradient. Clarified lysate was loaded onto an iodixanol gradient consisting of 15%, 25%, 40%, and 60% iodixanol phases prepared in PBS supplemented with NaCl and MgCl_2_ and centrifuged at 200,000 × g for 1 hour at 18°C in a Beckman Ti45 rotor (or equivalent). The 40% iodixanol fraction, which contains purified AAV particles, was collected in 1 mL fractions by side puncture with an 18-gauge needle. An SDS-PAGE was ran on each fraction to determine purity. Pure fractions were combined in a 100 kDa MWCO centrifugal filter unit (Amicon Ultra, Millipore) for buffer exchange into PBS and concentration. Purified vectors were stored at −80°C in single-use aliquots.

Vector genome titers were determined by droplet digital PCR (ddPCR) using primers and a probe targeting the GFP transgene or the AAV ITR sequence. Reactions were performed according to the manufacturer’s protocol (Bio-Rad QX200). Vectors were produced at titers of 10^12^ viral genomes per milliliter (vg/mL).

### Lentivirus production

Lentiviral vectors were produced by transfection of HEK293T cells using a second-generation packaging system. HEK293T cells were seeded 24 hours prior to transfection and co-transfected with four plasmids: (1) the lentiviral transfer plasmid pSIN-CK8-GFP; (2) psPAX2 packaging plasmid, and (3) pMD2.G envelope plasmid were transfected with Lipofectamine 2000. At 16–18 hours post-transfection, the medium was replaced with fresh complete medium. Virus-containing supernatant was collected at 48 or 72 hours post-transfection, pooled, and filtered through a 0.45 μm low-protein-binding membrane to remove cellular debris.

Lentiviral particles were concentrated using 40% polyethylene glycol for 30 minutes and then spun at 1500 x g for 45 minutes. The pellet was resuspended in PBS or an appropriate storage buffer, aliquoted, flash frozen and stored at −80°C.

### Bulk RNA-sequencing

RNA samples were collected from untreated, AAV6-treated, and lentivirus-treated iPSC-derived cardiomyocytes at Day 20 and Day 30 of differentiation. Cells were lysed using the RNeasy Mini Kit (Qiagen) according to the manufacturer’s protocol, including an on-column DNase I digestion step to eliminate genomic DNA contamination. Samples were sent to Novogene for quality control and bulk RNA-seq with poly(a) mRNA enrichment. For downstream analysis, sequencing reads were quasi-mapped, then counted to human GRCh38.p14 using Salmon (version 1.8.0) with default settings for gene- and transcript-level quantifications. Salmon results were read into R, using the package tximport. Differential gene expression analysis was performed on raw counts, using the R package DESeq2. Genes and transcripts with more than 100 raw counts were considered as expressed. PCA and hierarchical clustering were performed based on transcript-specific quantification. Variance-stabilization transformed data from DESeq2 were used as the input for PCA and hierarchical clustering. To define DEGs, cutoff values of absolute log2 fold change greater than 1 and adjusted p value less than 0.05 were used. DEGs were determined by performing pairwise comparisons across all treatment groups and time points. Z scores were used to generate heatmaps, using the R package Complex Heatmap.

## ACKNOWLEDGMENTS

The authors thank Sue and Bill Gross Stem Cell Research Center Core Facility for their support as well as Chris Denning, PhD, for the human induced pluripotent stem cell line. This study was supported by grants from the National Institutes of Health (K01HL169413 and R03HL183114 to A.E.), the American Heart Association (https://doi.org/10.58275/AHA.24CDA1259424.pc.gr.193591) to A.E., the Muscular Dystrophy Association (https://doi.org/10.55762/pc.gr.157032) to A.E., and T32 Predoctoral Training Fellowship (T32AR083870) to A.R.K.

## Author Contributions

A.E. conceived the project. E.C.L. and A.E. designed the experiments. E.C.L. performed the experiments and collected the data. E.C.L. and A.R.K. analyzed the results. E.C.L. and A.E. wrote the manuscript. A.E. oversaw all aspects of this work. A.E. and A.R.K. wrote grant applications that funded this project.

## REFERENCES

1. Duan D. Systemic AAV Micro-dystrophin Gene Therapy for Duchenne Muscular Dystrophy. Mol Ther. 2018 Oct 3;26(10):2337–56. PubMed PMID: 30093306.

2. Canessa EH, Spathis R, Novak JS, Beedle A, Nagaraju K, Bello L, et al. Characterization of the dystrophin-associated protein complex by mass spectrometry. Mass Spectrom Rev. 2024;43(1):90–105. PubMed PMID: 36420714.

3. Houang EM, Sham YY, Bates FS, Metzger JM. Muscle membrane integrity in Duchenne muscular dystrophy: recent advances in copolymer-based muscle membrane stabilizers. Skelet Muscle. 2018 Oct 10;8:31. PubMed PMID: 30305165; PubMed Central PMCID: PMC6180502.

4. Kamdar F, Garry DJ. Dystrophin-Deficient Cardiomyopathy. J Am Coll Cardiol. 2016 May 31;67(21):2533–46. PubMed PMID: 27230049.

5. Bergmann O, Bhardwaj RD, Bernard S, Zdunek S, Barnabé-Heider F, Walsh S, et al. Evidence for cardiomyocyte renewal in humans. Science. 2009 Apr 3;324(5923):98–102. PubMed PMID: 19342590; PubMed Central PMCID: PMC2991140.

6. Research C for BE and. FDA Investigating Deaths Due to Acute Liver Failure in Non-ambulatory Duchenne Muscular Dystrophy Patients Following ELEVIDYS. FDA [Internet]. 2025 Jun 25 [cited 2026 May 27]. Available from: https://www.fda.gov/vaccines-blood-biologics/safety-availability-biologics/fda-investigating-deaths-due-acute-liver-failure-non-ambulatory-duchenne-muscular-dystrophy-patients

7. Duan D. Lethal immunotoxicity in high-dose systemic AAV therapy. Mol Ther. 2023 Nov 1;31(11):3123–6. PubMed PMID: 37822079; PubMed Central PMCID: PMC10638066.

8. Duan D, Herzog RW. Deaths in gene therapy of Duchenne muscular dystrophy and other diseases: Underlying mechanisms and mitigating strategies. Mol Ther J Am Soc Gene Ther. 2026 Apr 1;34(4):1893–908. PubMed PMID: 41502088; PubMed Central PMCID: PMC12882805.

9. Kaufman BD, Veerapandiyan A, Soslow JH, Wittlieb-Weber C, Esteso P, Olson AK, et al. Taking ACTION to detect myocarditis related to recombinant gene transfer therapy for Duchenne Muscular Dystrophy; Consensus recommendations for cardiac surveillance. J Neuromuscul Dis. 2025 Mar;12(2):173–82. PubMed PMID: 39973402; PubMed Central PMCID: PMC13142859.

10. Emami MR, Espinoza A, Young CS, Ma F, Farahat PK, Felgner PL, et al. Innate and adaptive AAV-mediated immune responses in a mouse model of Duchenne muscular dystrophy. Mol Ther Methods Clin Dev. 2023 Sep 14;30:90–102. PubMed PMID: 37746243; PubMed Central PMCID: PMC10512012.

11. Costa-Verdera H, Meneghini V, Fitzpatrick Z, Abou Alezz M, Fabyanic E, Huang X, et al. AAV vectors trigger DNA damage response-dependent pro-inflammatory signalling in human iPSC-derived CNS models and mouse brain. Nat Commun. 2025 Apr 18;16(1):3694. PubMed PMID: 40251179; PubMed Central PMCID: PMC12008376.

12. Eguchi A, Gonzalez AFGS, Torres-Bigio SI, Koleckar K, Birnbaum F, Zhang JZ, et al. TRF2 rescues telomere attrition and prolongs cell survival in Duchenne muscular dystrophy cardiomyocytes derived from human iPSCs. Proc Natl Acad Sci U S A. 2023 Feb 7;120(6):e2209967120. PubMed PMID: 36719921; PubMed Central PMCID: PMC9963063.

13. Keegan AR, Gonzalez AFGS, Zhang R, Doan H, Torres-Bigio SI, Lau F, et al. Microdystrophins partially rescue deficits of duchenne muscular dystrophy iPSC-cardiomyocytes. Mol Ther Adv. 2026 Jul 2;34(3):201801. PubMed PMID: 42494980; PubMed Central PMCID: PMC13393429.

14. Chang ACY, Pardon G, Chang ACH, Wu H, Ong SG, Eguchi A, et al. Increased tissue stiffness triggers contractile dysfunction and telomere shortening in dystrophic cardiomyocytes. Stem Cell Rep. 2021 May 20;16(9):2169–81. PubMed PMID: 34019816; PubMed Central PMCID: PMC8452491.

